# Dynamics of *Salmonella* inoculated during rearing of black soldier fly larvae (*Hermetia illucens*) on chicken feed

**DOI:** 10.1101/2021.04.13.439665

**Authors:** J. De Smet, D. Vandeweyer, L. Van Moll, D. Lachi, L. Van Campenhout

## Abstract

The black soldier fly is currently the most produced edible insect on industrial scale, with its larval stage being processed into animal feed as the main application. As this insect species enters the feed and food chain, good hygiene and monitoring practices are needed to avoid the entrance of foodborne pathogens via the larvae. However, insufficient data on the risk of such introductions via industrial larvae production are available. To address this gap, a range of rearing trials were conducted in which the substrate, chicken feed, was inoculated with different levels of *Salmonella* and in which total viable counts and *Salmonella* counts were determined during the following days. The outgrowth of *Salmonella* was slower in those experiments with a lower initial contamination level than in experiments with a higher level. No significant reducing effect originating from the larvae on the substrate *Salmonella* counts was observed, in contrast to previous studies using other substrates. Our study also revealed that airborne transmission of *Salmonella* is possible under rearing conditions corresponding to those applied at industrial production sites. Based on our results, we recommend insect producers to use substrate ingredients free of *Salmonella*, and not to count on the antimicrobial activities that BSFL may exert in some situations towards food pathogens. More inoculation studies using other *Salmonella* serotypes, other zoonotic bacteria, other substrates, larvae of other ages and including variations on rearing protocols are needed in order to obtain a general view on the dynamics of food pathogens in this insect species and to support comprehensive risk assessments.

## 1. Introduction

The mass production of insects is now widely accepted as an agricultural activity in the Western world. Depending on the insect species, they can be used in human food and animal feed as an alternative source for proteins, but they also deliver other components, such as lipids for biodiesel production and biochemicals for cosmetics. They can also be applied in waste processing to support circularity in the bio-economy (Sogari et al., 2019). Particularly when insects are processed into food or feed products, safety hazards have to be monitored during rearing and processing to ensure a safe end product (van der Fels-Klerx et al., 2018). The insect species currently produced in the largest volume is the black soldier fly (*Hermetia illucens*) and the major application of its larval stage is as animal feed ingredient (Arru et al., 2019).

In 2015, the European Food Safety Authority (EFSA) published an initial risk profile for the production and consumption of edible insects for food and feed and listed potential safety hazards (EFSA Scientific Committee, 2015). Specific attention was paid to microbiological hazards, including *Salmonella*. Later, studies focused on the microbial composition of the larvae during rearing and the presence of food pathogens. For example, the presence of *Salmonella* sp. was observed in the residue of a black soldier fly larvae (BSFL) rearing cycle at an industrial setting, though no *Salmonella* was found in 25 g samples of the larvae (Wynants et al., 2018). Hence, even when using only the food- and feed-grade substrates that are currently allowed for insect rearing, good hygiene and monitoring practices are needed to avoid the introduction of this and likely also other foodborne pathogens in the feed and food chain via insects.

In Europe, the use of processed BSFL, or so-called ‘insect meal’, is currently allowed in aquafeed. Authorization in poultry and pig feed is to be expected (Byrne, 2021), and then larvae will enter the feed chain at an even large scale. Hence, monitoring and surveillance programs will have to upscale concomitantly. Information on the killing effect on food pathogens present in the larvae of post-harvest processing is very scarce, even though the aim should be to rear pathogen-free larvae and to avoid any introduction of food pathogens in larvae processing plants. In addition, the legislation in Europe (Regulation (EU) No 2017/893) currently allows the feeding of live insects to poultry, which is shown to benefit poultry welfare (Ipema et al., 2020). *Salmonella* can asymptomatically colonize the small intestine of poultry, along with the cecum, and therefore broilers and layers belong to the most likely vectors for *Salmonella* transmission to humans via food consumption (Cosby et al., 2015). Finally, in its brochure called “Three research priorities”, the European insect federation IPIFF (International Platform of Insects for Food and Feed), the first priority mentioned is to explore substrates for insect rearing that are not yet allowed but can further boost the contribution of the sector to a circular economy. Examples of envisaged streams are former foodstuffs containing meat, slaughter waste, etcetera. It goes without saying that in these types of substrates, the surveillance and control of food pathogens such as *Salmonella* will be of utmost importance. All mentioned facts underpin the high need for more data on the dynamics of food pathogens in BSFL, and in particular in the situation when they enter the rearing cycle via the substrates fed to the insects.

A typical approach to study the behavior of a zoonotic pathogen during rearing of or bioconversion by insects, is to inoculate the substrate with the micro-organism, provide it to the insects and follow-up possible colonization of the substrate and insects via classical microbial counts and/or sequencing of the whole microbiota. Some studies were performed in this way for BSFL in combination with a few zoonotic pathogens. The larvae were reported to be able to reduce the load of *Escherichia coli, Salmonella* spp., and *Enterococcus* spp. in their substrate by even up to 8 log cycles in some cases (Erickson et al., 2004; Lalander et al., 2013; Lalander et al., 2015; Liu et al., 2008; Lopes et al., 2020). It must be mentioned, though, that the substrate in all aforementioned studies was some type of manure and in one study aquaculture waste, and that the main aim was to find out whether BSFL can be used as bioconversion and sanitizing step in the processing of the waste. These publications, together with an increasing number of reports on the detection and description of a wide range of antimicrobial peptides in BSFL and antimicrobial effects of extracts (Choi et al., 2012; Xu et al., 2020), can lead to the general impression that the presence of food pathogens in whatever substrate of BSFL is not a large risk, since their growth is expected to be suppressed by the larvae. Although BSFL indeed may exert antimicrobial activity against specific micro-organisms and in certain conditions, more data are needed to elucidate the consequences of the presence of food pathogens in the substrate, especially when the larvae are produced as feed ingredient. The production of antimicrobial peptides has proven to be diet-dependent (Vogel et al., 2018), and so may be the possible pathogen reducing effect. Even for substrates currently allowed and frequently used in industrial BSFL production for animal feed, the interactions between a pathogen such as *Salmonella*, the larvae and the other micro-organisms present during rearing are not yet uncovered. It is not known for allowed substrates, if and how fast *Salmonella* can colonize the substrate and/or the larvae, and which factors, such as the contamination level of the pathogen, the type of substrate and other rearing conditions, the background microbiota and the overall hygiene level of the production environment, the age of the larvae, the *Salmonella* serotype(s) present, influence the interactions.

The aim of this work was to conduct a range of rearing trials with BSFL after inoculating the substrate with *Salmonella* and to determine total viable counts and *Salmonella* counts in the days after providing the inoculated substrate. While there are, as mentioned before, many factors that possibly affect the dynamics of *Salmonella*, we opted to perform all inoculation experiments with the same substrate and in the same rearing conditions. A factor that was varied, however, was the contamination level. The research was started first with trials at a high contamination level, to mimic worst-case scenarios, and then moved on to lower levels, probably implying more realistic scenarios in industry. As substrate chicken feed was chosen, which is frequently used in research as well as in (the first stages of rearing in) industry. The chicken feed was not frozen or autoclaved, so that the endogenous microbiota of the feed was also active during the experiments.

## 2. Material and methods

### 2.1. Overview of consecutive series of experiments

To study the behavior of *Salmonella* during BSFL rearing, three different experimental set-ups, conducted at laboratory scale, were used. In a first series of experiments, wild-type *Salmonella* strains were used for inoculation. Since this resulted in the presence of a large quantity of non-specific colonies (background microbiota) on the selective plates (as discussed in detail in the results), kanamycin resistant mutants were used in the second series, and two inoculation levels were included. In this second series, airborne contamination between the conditions tested (as described further) could not be excluded. Therefore, a third series of experiments was conducted in which the conditions were incubated separately. All experimental set-ups were performed with two or three separate batches of larvae (meaning that the larvae were reared independently during different rearing cycles), and for each repetition, two replicates (i.e. containers with larvae) were included. A general overview of the experimental designs and the varied parameters can be found in Table 1. For each experiment, four different rearing conditions were included, an overview of which is given in Figure 1: (i) substrate without *Salmonella* and without larvae (S), (ii) substrate inoculated with pathogen *Salmonella* but without larvae (S+P), (iii) substrate with larvae but without *Salmonella* (S+L), and (iv) substrate inoculated with *Salmonella* and provided with larvae (S+P+L).

**Table 1:** Overview of the consecutive series of experiments.

| Series of experiments | *Salmonella* strains included | Target level for inoculation | Incubation | Number of batches (and of replicates) |
| --- | --- | --- | --- | --- |
| 1 | *S. Typhimurium*, *S. Infantis*, *S. Enteritidis* | 7 log CFU/g | Inoculated and Uninoculated conditions incubated together | 2 (2) |
| 2 | *S. Typhimurium* KAN<sup>R</sup><br>*S. Infantis* KAN<sup>R</sup> | 4 log CFU/g<br>7 log CFU/g | Inoculated and Uninoculated conditions incubated together | 2 (2) |
| 3 | *S. Typhimurium* KAN<sup>R</sup><br>*S. Infantis* KAN<sup>R</sup> | 3 log CFU/g | Inoculated and Uninoculated conditions incubated separately | 3 (2) |

**Figure 1:**
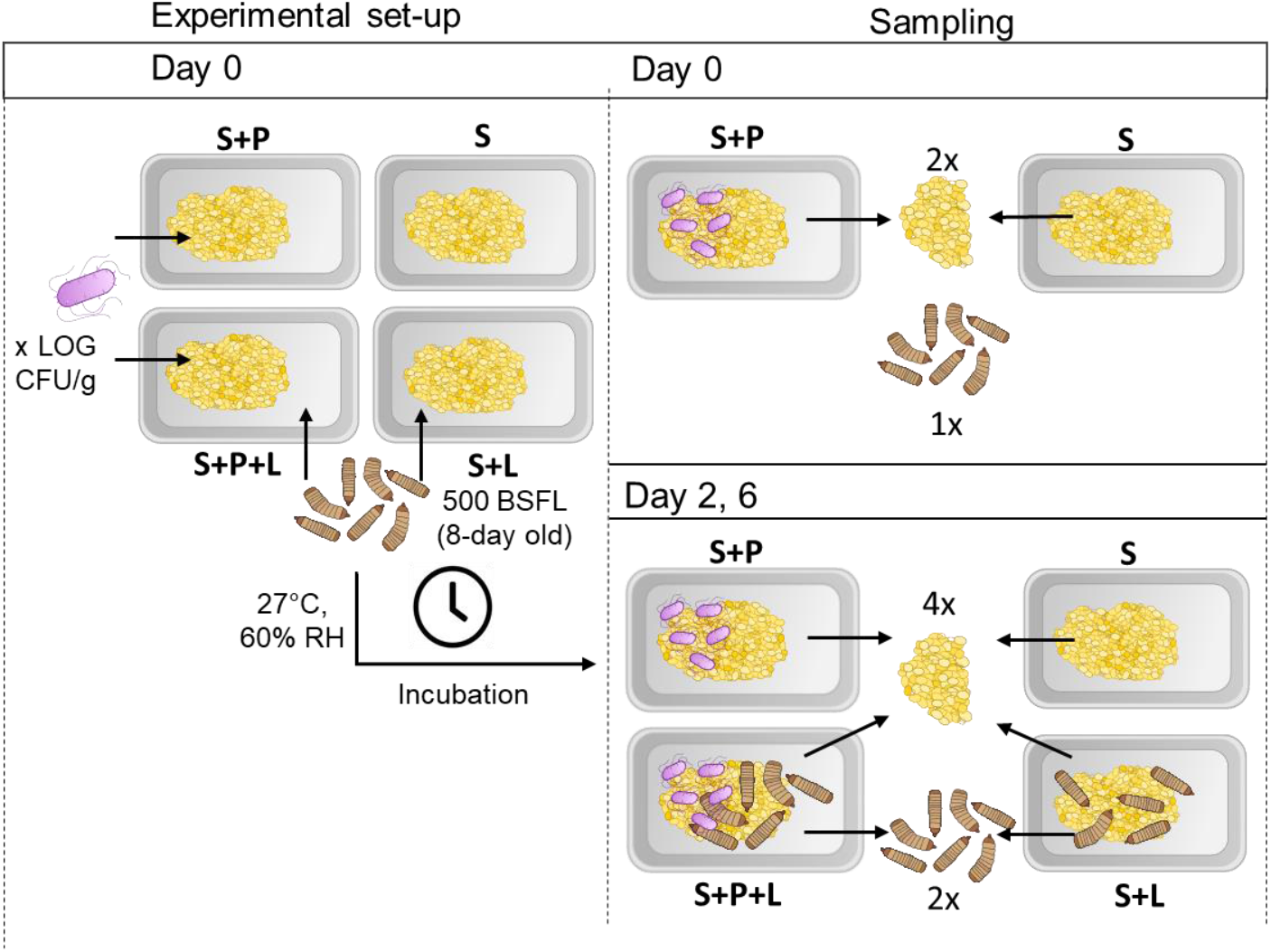
Schematic overview of experimental set-up and sampling method in a challenge experiment, depicted for one batch of larvae. S = substrate without *Salmonella* and without larvae, S+P = substrate inoculated with pathogen *Salmonella* but without larvae, S+L substrate with larvae but without *Salmonella*, and S+P+L = substrate inoculated with *Salmonella* and provided with larvae.

### 2.2. Salmonella cultivation and creation of kanamycin resistant strains

Three different *Salmonella* strains were used: *Salmonella enterica* subsp. *enterica* serovar Enteritidis (LMG 18735), *Salmonella enterica* subsp. *enterica* serovar Typhimurium (LMG 18732) and *Salmonella enterica* subsp. *enterica* serovar Infantis (LMG 18746), all purchased from the Belgian Coordinated Collection of Microorganisms (BCCM). For experiment series 1, a mixture of all three bacterial strains was used for inoculation. From design 2 onwards, a mixture of kanamycin resistant *S. typhymurium* and kanamycin resistant *S. infantis* was used.

Antibiotic resistant strains were generated by using a temperature sensitive pHSG415-tnsABCD helper plasmid and a modified mini-Tn7 delivery system as described previously (Shivak et al., 2016). In our study, the delivery plasmid pUC18R6K-mini-Tn7T-pCS26-KmR sig70_c10 LUX was used to incorporate a kanamycin resistance gene into the target bacteria. To achieve chromosomal integration and kanamycin resistance, electrocompetent *Salmonella* cells were made, as described by Shivak et al. (2016). Next, both the helper plasmid and delivery plasmid were transformed via electroporation. After electroporation, cells were allowed to recover in SOC medium (2% tryptone (Lab M, UK), 0.5% yeast extract (VWR, Belgium), 10 mM NaCl (Acros Organics, Belgium), 2.5 mM KCl (Acros Organics), 10 mM MgCl2 (Acros Organics), 10 mM MgSO4 (Acros Organics), and 20 mM glucose (Acros Organics)) for 2 h at 28°C. Then, the transformation mix was plated on LB/Kan^50^ (Luria Bertani, composed of 10.0 g/l peptone (Biokar Diagnostics, France), 5.0 g/l yeast extract, 5.0 g/l NaCl, 15 g/l agar (VWR, Belgium), 50 μg/ml Kanamycine (Thermo Fisher Scientific, Belgium)) and incubated overnight at 37°C. The correct chromosomal integration of the resistance cassette was checked in the obtained transformants with PCR using two primer pairs (primer pair 1: glmSdetectFor - lux-check and primer pair 2: glmSdetectRev-KmCheck). The conditions for the PCR reaction can be found in Supplementary table 1.

### 2.3. Inoculation of substrate

The selected *Salmonella* species were grown overnight at 37°C in Luria Bertani broth (see LB plates but without agar), and with Kan^50^ if resistant strains were used (design 2 and 3). Then, the overnight cultures were diluted using LB to a McFarland unit (MFU, DEN-1 McFarland Densitometer, Grant instruments, UK) of 5. Next, the different *Salmonella* strains were combined in equal volumes to create a suitable *Salmonella* mixture. For the substrate, chicken starter feed (Startmeel voor Kuikens 259, AVEVE, Belgium) was grinded with a mixer (Espressions EP9800 Powerblender) using two times the “Ice Crush” program. The substrate was then prepared by mixing the grinded chicken starter feed and tap water in a 1:1 ratio (w/v), and 100 g of wetted feed was placed in polypropylene trays (1L). Finally, for all conditions which required a challenge (S+P and S+P+L), an aliquot of the prepared inoculum was added to 100 g of wetted feed to obtain a desired starting concentration of 7 log CFU/g (3.3 ml of a MFU 5 solution) (design 1) or 7 and 4 (1.2 ml of a 1/2000 diluted MFU 2 solution) log CFU/g (design 2) or 3 log CFU/g (0.6 ml of a 1/2000 diluted MFU 2 solution) (design 3). The inoculated feed was homogenized using a sterile spoon. To the uninoculated groups, an volume of sterile LB broth was added equal to their inoculated counterparts.

### 2.4. Rearing of BSFL and sampling

For this study, BSFL were supplied by and originated from a colony maintained by RADIUS (Thomas More University University of Applied Sciences, Geel, Belgium). They were nursed until day 8 on a mixture of chicken starter feed and tap water in a 1:1 ratio (w/w) in a climate chamber (Pharma 600, Weiss Technik, Belgium) at 27°C and 60% relative humidity (RH). At that time, the challenge experiment was initiated. Approximately 500 (as determined by the average weight of three times 10 larvae) 8-day-old BSFL were added to the container of each larvae-containing replicate (S+L and S+P+L). The dimensions of the containers used were 10 cm x 15 cm, yielding a density of 3.3 larvae/cm^2^. The containers were fitted with a lid containing a mesh covered surface (7 cm x 13 cm) to allow air circulation, but prevent larval escape. Next, 100 g of either or not contaminated feed was added to the respective replicates and all containers were placed in a climate chamber (Memmert HPP260, Memmert, Belgium) at 27°C and 60% relative humidity (RH) until the end of the experiment, which was 6 days after the challenge. On day 2 and day 4, an additional 80 g of uncontaminated feed (chicken starter and tap water) was added to each rearing box. For design 1 and 2, all experimental conditions were incubated together in the same incubator. For design 3, replicates with different conditions were incubated separately to avoid cross-contamination.

Sampling of larvae and substrate took place on day 0, day 2 and day 6 in aseptic conditions (Figure 1). Larvae were separated from the substrate by sieving. Then they were disinfected prior to sampling in three subsequent washing steps: a first disinfection step with 100 mL of 70% ethanol (1 minute at 200 rpm on a laboratory shaking table (Unimax 1010, Germany)) was followed by two rinsing steps with 100 mL of sterile, demineralized water (1 minute each at 200 rpm on the shaking table). To monitor the growth of the larvae, the mass of 10 larvae was measured and this was repeated 5 times per replicate. Next, 5 g of larvae were collected and diluted tenfold in physiological peptone solution (PPS, 0.85% NaCl, 0.01% peptone) before pulverizing the larvae in the solution by using an ethanol sterilized home type mixer (Bosch CNHR 25). Similarly, 5 g of substrate was sampled from each replicate and diluted 1:10 in PPS. Prior to microbiological analysis, both larval and substrate samples were also homogenized using a stomacher (BagMixer 400CC, Interscience, France) for 1 minute.

### 2.5. Microbiological analysis

For each sample, the total aerobic viable count and the specific *Salmonella* count were determined. All plate counts were performed according to the ISO standards for microbial analyses of food and feed as compiled by Dijk et al. (2015). For the total aerobic viable count, serial dilutions were made in PPS, plated on Plate Count Agar (PCA, Biokar Diagnostics, Beauvais, France), and incubated at 30°C for 72 hours. *Salmonella* was counted by plating the diluted samples on a chromogenic RAPID’ *Salmonella* agar (BioRad Laboratories, Belgium) and incubating the plates at 37°C for 24 hours. The cell density of all inocula was also verified by plating a serial dilution on both the RAPID’ *Salmonella* agar and PCA.

### 2.6. Statistical analysis

For design 1 to 3, total viable counts as well as *Salmonella* sp. counts of larvae and substrate samples for each condition were compared between sampling moments using one-way ANOVA, with Tukey HSD as post hoc test in case of equal variances. When the variances were not equal, the Welch’s ANOVA with Steel–Dwass All Pairs post hoc test was used. Furthermore specific pair-wise t-tests were conducted in order to compare total viable counts and/or *Salmonella* sp. counts between samples at day 6. When counts of a sample were below the detection limit, the detection limit itself was chosen as value to be included for statistical analysis. All these tests were performed using JMP Pro 15.0.0 from SAS. For each test, a significance level of α = 0.05 was considered.

## 3. Results

### 3.1. Salmonella dynamics using a high inoculation level

The first goal was to examine a possible worst-case scenario during rearing. This was achieved in experimental design 1, by contaminating the substrate and aiming at an inoculation level of 7 log CFU of *Salmonella* sp. per g substrate. The actual *Salmonella* sp. counts reached at day 0 were close to that target level, being 7.4 ± 0.5 log CFU/g (Table 2). No impact on the larval growth was observed from the presence of *Salmonella* sp. in the substrate (Supplementary figure 1), which was to be expected, as no evidence is present in literature that *Salmonella* sp. would be pathogenic for BSFL (Joosten et al., 2020).

**Table 2:** Total aerobic viable counts and *Salmonella* spp. counts of larvae and substrate samples of experiment series 1, involving a high contamination of wild-type *Salmonella* strains (7 log CFU/g) and all conditions incubated in the same climate chamber. Values present the mean (± standard deviation) of two batches, each with two replicates per condition (n=2×2). S = Substrate; S+P = Substrate with pathogen; S+L = Substrate with larvae; and S+P+L = Substrate with pathogen and larvae.

| Experimental condition | Investigated sample | Target *Salmonella* spp. contamination level in substrate (log CFU/g) | Total viable count (log CFU/g) |  |  | *Salmonella* sp. count (log CFU/g) |  |  |
| --- | --- | --- | --- | --- | --- | --- | --- | --- |
|  |  |  | Day 0 | Day 2 | Day 6 | Day 0 | Day 2 | Day 6 |
| S | substrate | 0* | $6.5 \pm 2.3^a$ | $11.1 \pm 0.9^a$ | $11.6 \pm 0.3^a$ | $<2.0 \pm 0.0^{\dagger,a}$ | $<2.0 \pm 0.1^{\dagger,a}$ | $<2.0 \pm 0.0^{\dagger,a}$ |
| S + P | substrate | 7 | $8.0 \pm 0.3^a$ | $11.4 \pm 0.5^a$ | $12.1 \pm 1.0^a$ | $7.4 \pm 0.5^a$ | $8.1 \pm 0.4^a$ | $9.5 \pm 0.8^b$ |
| S + L | substrate | 0* | $6.5 \pm 2.3^{\circ,a}$ | $>11.5 \pm 1.2^a$ | $12.4 \pm 1.0^a$ | $<2.0 \pm 0.0^{\circ,\dagger,a}$ | $<3.3 \pm 2.6^{\ddagger,a}$ | $<6.3 \pm 5.0^{\ddagger,a}$ |
| S + P + L | substrate | 7 | $8.0 \pm 0.3^{\circ,a}$ | $>11.0 \pm 1.7^a$ | $11.9 \pm 0.7^a$ | $7.4 \pm 0.5^{\circ,a}$ | $7.9 \pm 1.7^a$ | $8.3 \pm 1.7^a$ |
| S + L | larvae | 0* | $9.1 \pm 0.5^a$ | $10.3 \pm 0.8^b$ | $10.7 \pm 0.5^b$ | $<2.0 \pm 0.0^{\dagger,a}$ | $<3.7 \pm 2.0^{\ddagger,a}$ | $<5.7 \pm 4.2^{\ddagger,a}$ |
| S + P + L | larvae | 7 | | $11.0 \pm 1.0^a$ | $10.5 \pm 0.3^a$ | | $6.5 \pm 1.2^a$ | $7.3 \pm 1.0^a$ |

\*Uninoculated replicates;
$^{\circ}$ = sample is similar at this timepoint to chicken starter feed without larvae and was not determined a second time;
$^{\dagger}$ "<2.0" indicates that *Salmonella* sp. was below the detection limit (2 log CFU/g) in every sample;
$^{\ddagger}$ "<" followed by a value higher than 2.0 log CFU/g indicates that *Salmonella* sp. was below the detection limit in at least one, but not all samples.
<sup>abc</sup> Average values for total viable counts and *Salmonella* sp. counts within each row that share a letter in superscript did not significantly ( $p \geq 0.05$ ) increase or decrease between sampling days.

A clear increase was observed in the total aerobic viable count from day 0 till day 6 for all substrate samples, irrespective of the presence of larvae, with counts increasing from 6.5-8.0 log CFU/g to 11.6-12.4 log CFU/g (Table 2). For the substrate without larvae (S and S+P), this is in contrast to observations made in earlier challenge experiments reported for *Tenebrio molitor*, where the total viable counts did not significantly change over a same period of time for that type of sample (Wynants et al., 2019). A plausible explanation is that the higher moisture content of the substrate for BSFL (approximately 50%) is more suited for microbial growth compared to that of the substrate for *Tenebrio molitor*, that consisted of dry wheat bran with carrot pieces as moisture source. Another observation is that the presence of larvae does not seem to influence the microbial load (at least, in quantitative terms) of the substrate in any way (Table 2).

At day 0, the total aerobic viable count of the larvae (in S+L as well as in S+P+L) was higher than that of the substrate, and with 9.1 ± 0.5 log CFU/g, this number is in the range of earlier reports on the total viable count present in larvae (Wynants et al., 2018). The increase over time in their interior microbial load was also much less pronounced than the increase in their substrate. Final numbers, ranging from 10.5 to 10.7 log CFU/g, remained approximately one log CFU lower than the microbial load in the substrate. This likely can be explained by the larval interior being completely occupied as ecological niche, in terms of nutrient availability and/or it can point towards the presence of control mechanisms restricting the microbial load inside the larvae.

From the *Salmonella* sp. counts, it is clear that the pathogen can thrive well in the substrate as their number increased significantly in the inoculated, larvae-free condition (S+P) from 7.4 ± 0.5 log CFU/g at day 0 to 9.5 ± 0.8 log CFU/g at day 6 (Table 2). At the same time, *Salmonella* was not detected (detection limit is 2 log CFU/g) in the uninoculated substrate without larvae (S) at day 0, 2 and 6. Interestingly, this was different in the uninoculated substrate samples with larvae (S+L). Here, *Salmonella* was detected from day 2 in two out of four replicates and its counts increased further in these replicates to reach counts close to those of the substrates of the inoculated experiments (S+P and S+P+L, Table 2). Two explanations are possible here. First, all larval samples showed *Salmonella* levels below the detection limit at day 0, but it cannot be excluded that it was present below the detection limit and started growing to detectable levels during the test. Secondly, and according to us more likely, cross-contamination occurred in the climate chamber due to airborne transmission of *Salmonella*, as will be further addressed in the discussion. In contrast to the studies mentioned in the Introduction Section, no significant decreases in *Salmonella* sp. counts were observed over time in the presence of larvae, neither in the substrate samples nor in the larval samples (Table 2). Overall, the *Salmonella* counts in the larvae were approximately between 1 and 2 log CFU/g lower than in the corresponding substrate sample.

The aforementioned observations were hindered by a background growth of micro-organisms on the selective medium for *Salmonella* count (Supplementary Figure 2). Though distinction with *Salmonella* and proper counting of *Salmonella* was still possible due to the colony morphology, and in particular the color, the large abundances of the background microbiota were unwanted. Indeed, the presence of the background microbiota could hinder the next step in our study to explore the impact of lower, more realistic, inoculation levels on the microbiological safety of BSFL. To circumvent this problem, all three *Salmonella* sp. used were genetically engineered to express a kanamycin resistance gene. The procedure was successful for *S*. Typhimurium and *S*. Infantis, so these two strains were mixed and used as inoculum in experimental design 2 and 3 (Table 1). The use of kanamycin, at a concentration of 50 μg/ml, indeed had a significant impact on the background growth (Supplementary Figure 2).

### 3.2. Salmonella dynamics using a high and low inoculation level and kanamycin-resistant strains

Using a mixture of the two resistant *Salmonella* strains, a second set of challenge experiments was executed (design 2). This design included both a low (4 log CFU/g), as well as the previous high (7 log CFU/g) inoculation level. No impact of the inoculations was observed on larval growth (data not shown), as in design 1. The microbiological results are shown in Table 3. The total viable counts were slightly lower over the whole design than the counts in design 1. However, the trends were comparable, showing a 4 to 5 log CFU/g increase in the substrate samples over the 6-day time frame. The initial total viable count in the uninoculated larvae (S+L) at day 0 was 8.2 ± 0.4 log CFU/g. This count only slightly increased to, on average, 9.2 ± 0.5 log CFU/g at day 6 and a similar observation was made for the inoculated larvae (S+P+L). As in design 1, the total viable count in the larvae was between 1 and 2 log CFU/g lower than the count reported in their corresponding substrates, which adds weight to the hypothesis that the ecological niche is fully occupied and/or that larvae control to some extent the total microbial load in their interior.

**Table 3:**
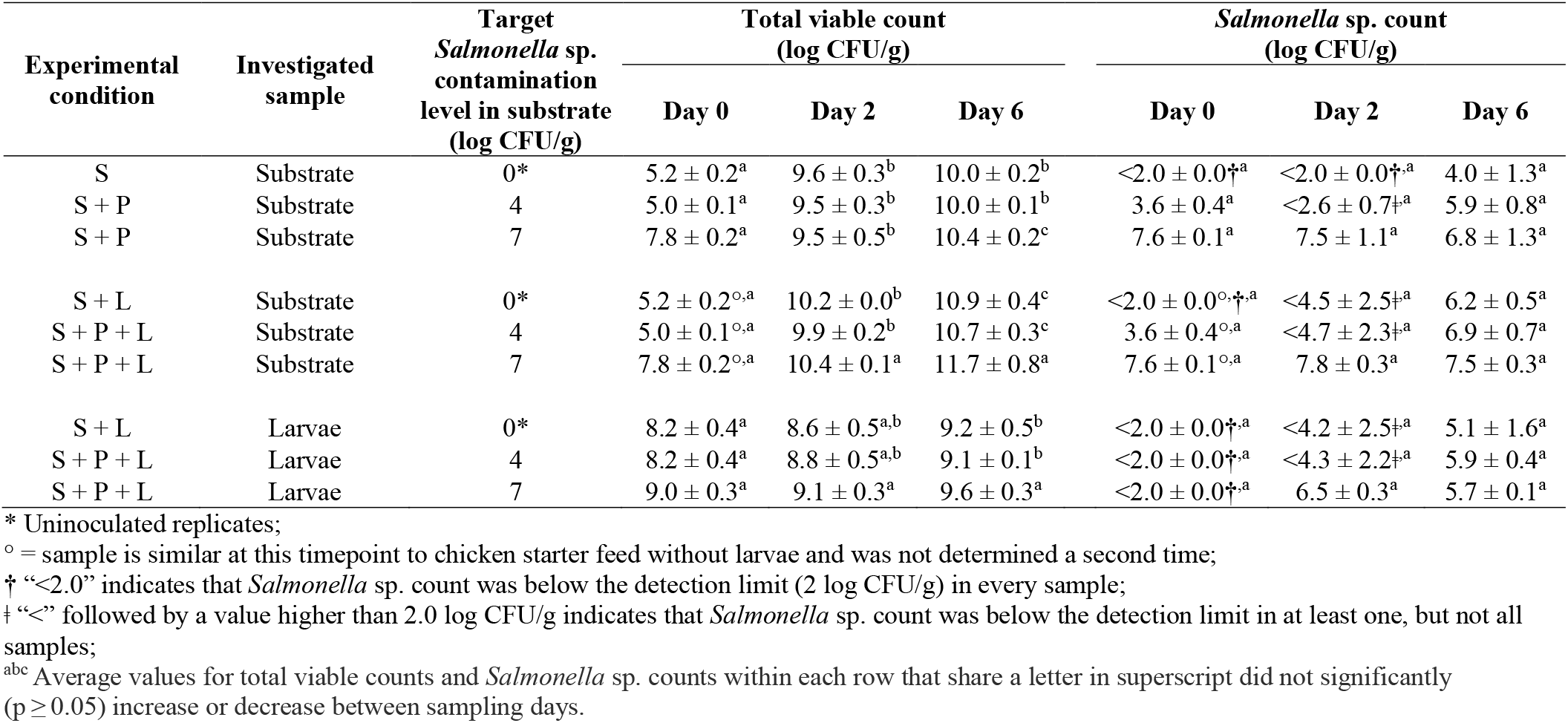
Total aerobic viable counts and *Salmonella* sp. counts of larvae and substrate samples of experiment series 2, involving both a low (4 log CFU/g) and high (7 log CFU/g) contamination of resistant *Salmonella* strains and all conditions incubated in the same climate chamber. Values represent the mean (± standard deviation) of two batches, each with two replicates per condition (n=2×2). S = Substrate; S+P = Substrate with pathogen; S+L = Substrate with larvae; and S+P+L = Substrate with pathogen and larvae.

For *Salmonella* (Table 3), the counts in the substrate (S+P) at day 0 were respectively 3.6 ± 0.4 log CFU/g and 7.6 ± 0.1 for the log 4 and log 7 challenge. At the high inoculation level, *Salmonella* counts remained fairly constant over time in the substrate, both in the absence (S+P) and presence (S+P+L) of larvae. In contrast, at the low inoculation level, an increase was observed over the 6-day period to a count of 5.9 ± 0.8 log CFU/g and 6.9 ± 0.7 log CFU/g in the absence (S+P) or presence (S+P+L) of larvae respectively. At day 2, considerable variation between replicates was observed in the substrate for both conditions (S+P and S+P+L), with *Salmonella* counts even below the detection limit in respectively two and one out of the four replicates. This indicates that at lower inoculation levels, the initial colonization speed of the substrate by *Salmonella* sp. is more variable and can even lead to an apparent removal of this pathogen. Nevertheless, the pathogen managed to successfully colonize the substrate over time. This observation is not impacted by the presence of larvae, as a similar trend is observed in both conditions and S+P and S+P+L at the 4 log CFU/g inoculation level.

Focusing on the larvae, for the 7 log CFU/g challenge test we observed a colonization evolution to 5.7 ± 0.1 CFU/g at day 6, which was a lower end value than in series 1 where a *Salmonella* count of 7.3 ± 1.0 was reached. Yet a similarity with series 1 was that the counts in the larvae both at day 2 and day 6 were about 1.5 log CFU/g and significantly (p < 0.001) lower than in the corresponding substrates. It should not be interpreted as a specific reduction of *Salmonella* numbers by the larvae during the rearing process, because this difference between larvae and their substrate is also observed for the total viable count. Moreover, after a challenge with 4 log CFU/g, at day 6 *Salmonella* reached even the same count (5.9 ± 0.4 CFU/g) as after the 7 log CFU/g challenge.

Extreme care that was taken in all series of experiments to avoid cross-contamination during manipulations of containers and samples. Therefore, the most intriguing observation from experiment series 2 was the presence of the *Salmonella* in the various uninoculated samples (S and S+L). Since kanamycin was used in the medium, the colonies found on the plates originated from resistant strains. All four replicates of the uninoculated samples (S+L), both in the substrate and larvae, were contaminated at day 6. With respectively 6.2 ± 0.5 and 5.1 ± 1.6 log CFU/g, these counts are comparable to the counts observed in the inoculated samples. All four replicates of the substrate (S) were also contaminated at day 6, while these replicates had a count below the detection limit at day 2. A possible hypothesis for these contaminations is that a small amount of naturally resistant *Salmonella* sp. were present below the detection limit and reached sufficient levels to be detected over time. Another possible explanation is the occurrence of airborne transmission of *Salmonella* in the climate chamber, which will be explored in more detail in the discussion. Though additional research is needed to confirm the following finding, the larvae seem to aggravate this transmission, as contaminations manifested themselves earlier if larvae were present (three out of four replicates of substrate contaminated at day 2 for S+L) compared to substrate without larvae, and the contaminations also reached a higher level in presence of larvae. Their movement through the substrate via their feeding behavior might contribute to airborne distribution via aerosols and/or dust particles.

### 3.3. Salmonella dynamics using a low inoculation level, kanamycin-resistant strains and prevention of airborne transmission

A third series of experiments with the same resistant strains was performed, and conditions were incubated separately from each other (i.e. consecutively), so that airborne transmission between the different conditions was completely excluded. An even lower contamination level than in the previous series of tests was applied, i.e. a target level of 3 log CFU/g *Salmonella*, to cover more potential situations in the insect industry related to *Salmonella*-infected substrates. The microbiological counts can be found in Table 4. As for the previous two series of tests, no impact on larval growth was observed (data not shown). The data for the total viable counts were similar to those found in the previous series, with the total viable counts increasing in all substrates, regardless of larval presence, to levels over 10 log CFU/g. At the same time, the total viable counts showed a smaller increase in the larvae than in the substrate and reached a level that was about 1 log CFU/g lower than that of the substrate.

**Table 4:**
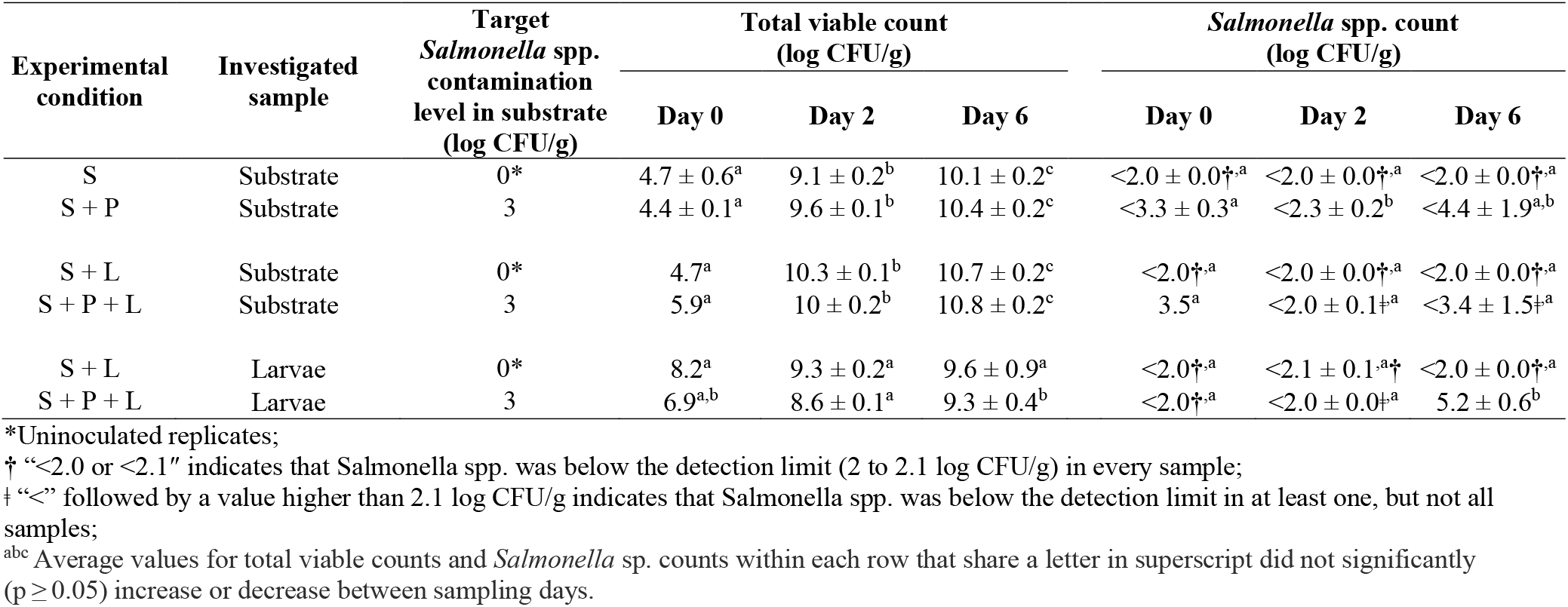
Total aerobic viable counts and *Salmonella* spp. counts from larvae and substrate samples of experiment series 3 involving a low contamination of resistant *Salmonella* strains (3 log CFU/g) and all conditions incubated separately in climate chamber. Values represent the mean (± standard deviation) of three batches, each with two replicates per condition (n=3×2). S = Substrate; S+P = Substrate with pathogen; S+L = Substrate with larvae; and S+P+L = Substrate with pathogen and larvae.

| Experimental condition | Investigated sample | Target *Salmonella* spp. contamination level in substrate (log CFU/g) | Total viable count (log CFU/g) |  |  | *Salmonella* spp. count (log CFU/g) |  |  |
| --- | --- | --- | --- | --- | --- | --- | --- | --- |
|  |  |  | Day 0 | Day 2 | Day 6 | Day 0 | Day 2 | Day 6 |
| S | Substrate | 0* | 4.7 ± 0.6<sup>a</sup> | 9.1 ± 0.2<sup>b</sup> | 10.1 ± 0.2<sup>c</sup> | <2.0 ± 0.0<sup>†,a</sup> | <2.0 ± 0.0<sup>†,a</sup> | <2.0 ± 0.0<sup>†,a</sup> |
| S + P | Substrate | 3 | 4.4 ± 0.1<sup>a</sup> | 9.6 ± 0.1<sup>b</sup> | 10.4 ± 0.2<sup>c</sup> | <3.3 ± 0.3<sup>a</sup> | <2.3 ± 0.2<sup>b</sup> | <4.4 ± 1.9<sup>a,b</sup> |
| S + L | Substrate | 0* | 4.7<sup>a</sup> | 10.3 ± 0.1<sup>b</sup> | 10.7 ± 0.2<sup>c</sup> | <2.0<sup>†,a</sup> | <2.0 ± 0.0<sup>†,a</sup> | <2.0 ± 0.0<sup>†,a</sup> |
| S + P + L | Substrate | 3 | 5.9<sup>a</sup> | 10 ± 0.2<sup>b</sup> | 10.8 ± 0.2<sup>c</sup> | 3.5<sup>a</sup> | <2.0 ± 0.1<sup>‡,a</sup> | <3.4 ± 1.5<sup>‡,a</sup> |
| S + L | Larvae | 0* | 8.2<sup>a</sup> | 9.3 ± 0.2<sup>a</sup> | 9.6 ± 0.9<sup>a</sup> | <2.0<sup>†,a</sup> | <2.1 ± 0.1<sup>a,†</sup> | <2.0 ± 0.0<sup>†,a</sup> |
| S + P + L | Larvae | 3 | 6.9<sup>a,b</sup> | 8.6 ± 0.1<sup>a</sup> | 9.3 ± 0.4<sup>b</sup> | <2.0<sup>†,a</sup> | <2.0 ± 0.0<sup>‡,a</sup> | 5.2 ± 0.6<sup>b</sup> |

\*Uninoculated replicates;
† “<2.0 or <2.1” indicates that *Salmonella* spp. was below the detection limit (2 to 2.1 log CFU/g) in every sample;
‡ “<” followed by a value higher than 2.1 log CFU/g indicates that *Salmonella* spp. was below the detection limit in at least one, but not all samples;
<sup>abc</sup> Average values for total viable counts and *Salmonella* sp. counts within each row that share a letter in superscript did not significantly ( $p \geq 0.05$ ) increase or decrease between sampling days.

No *Salmonella* was detected over the course of the challenge test in any of the uninoculated samples (S and S+L). In this third series of experiments, it can be excluded that *Salmonella* was present in the larvae under the detection limit at the start of the trial. In the same way as the 4 log CFU/g inoculation in series 2, for the inoculated substrate (S+P) a reduction occurred at day 2 to a level of < 2.3 ± 0.2 log CFU/g (with two out of six replicates below the detection limit). At day 6, the *Salmonella* counts were higher again, as they reached a value of < 4.4 ± 1.9 log CFU/g. Interestingly, in this design the two replicates that were below the detection limit at day 2 remained below the limit at day 6. In addition, at this low inoculation level, the larvae seemed to hinder colonization, since all six replicates of the substrate with pathogen and larvae (S+P+L) were below the detection limit at day 2. At day 6, four of the six replicates had a *Salmonella* count below detection, leading to an average count of < 3.4 ± 1.5 log CFU/g. This means that no significant increase in counts occurred in the substrate. This was not the case in the larvae, however. While no *Salmonella* was detected in the larvae on day 2, all replicates had detectable counts on day 6 with an average of 5.2 ± 0.6 log CFU/g. This number does not differ much from the final counts observed in experiment series 2 (Table 3) for the larvae at either of the two inoculation levels, indicating that the intestines of the larvae offer a habitable niche for *Salmonella* to colonize the larvae to a certain extent. That niche might even be more suited for *Salmonella* growth than the surrounding substrate, when only a low amount of *Salmonella* cells is present. The latter phenomenon might be substrate dependent, however, and needs further investigation.

## 4. Discussion

This work comprises inoculation experiments with the focus on the contamination level. In the industrial practice of BSFL rearing for feed purposes, *Salmonella* can be present in ingredients used to prepare the substrate mixture. Typical (and currently allowed) ingredients include cereals and cereal-based materials such as distillers dried grains with solubles, fruit and vegetables and derived products, former foodstuffs (purely vegetal or containing eggs or milk) and compound feed. Evidence is present in literature for each of these ingredient categories that they can contain *Salmonella* (Berghofer et al., 2003; Centers for Disease Control and Prevention (CDC), 1998; Gosling et al., 2021; Jongen, 2005; Lee et al., 2016; Eglezos, 2010). The contamination level of *Salmonella* in a substrate mixture ready to provide to BSFL can vary, because (1) the load in individual contaminated ingredients can vary, and (2) the ingredients and/or the finished feed may be stored before feeding to the insects, causing *Salmonella* counts to increase or reduce during storage. Since the *Salmonella* load of the substrate can vary and since it is not known whether or this load determines the fate of the pathogen in contact with larvae in their substrate, several inoculation levels were included in this research. While a lower inoculation level indeed pointed at a slower colonization by *Salmonella*, or a longer suppression of *Salmonella* by the larvae, in none of the cases investigated, the larvae were able to eradicate the pathogen completely, or even to reduce its counts over time. In trials with inoculated substrate and larvae (S+P+L), both substrate and larvae were still contaminated after 6 days and *Salmonella* counts were as high as or sometimes several log cycles higher than values at day 0. These results are in strong contrast with the reports mentioned earlier (Erickson et al., 2004; Lalander et al., 2013; Lalander et al., 2015; Liu et al., 2008; Lopes et al., 2020), that generally describe a pronounced reduction of *Salmonella* (and some other zoonotic bacteria) by BSFL treatment of the waste, investigated as a possible hygienization step. Several possible explanations can be put forward for this difference. The manure types and aquaculture waste can be expected to substantially differ from the substrate used in our study, both in terms of chemical and microbiological composition. *Salmonella* cells inoculated in manure or aquaculture waste may be faced with another and potentially less favorable nutrient profile and a larger background microbiota than the chicken feed in our study. Therefore, in this potentially tougher environment, the cells may have less competitive advantage and/or may be less fit, and in this way they may be more vulnerable for the antimicrobial mechanisms exposed by BSFL, such as antimicrobial peptides, lysozymes, other antimicrobial components and the low pH encountered during intestinal passage (Bonelli et al., 2019). The fact that the substrate is different, can also have an impact on the larvae, which may be more triggered by the chemical and microbiological composition of manure or aquaculture waste than that of chicken feed to activate their immune system and exert their antimicrobial activities (Vogel et al., 2017).

A difficulty encountered in our inoculation experiments was the fact that, when trying to specifically count the *Salmonella* inoculated in the experiments, colonies from the background microbiota were also present on the selective plates. This indicates that micro-organisms with properties very close to the pathogen are present in the background microbiota. In our study, we eliminated these organisms by using antimicrobial resistance as an additional selection mechanism in the plates. In our previous work on inoculation experiments with *Salmonella* during yellow mealworm rearing (Wynants et al., 2019a), no substantial background microbiota hindered the *Salmonella* counts, and an additional selection mechanism was therefore not necessary. The other studies on BSFL already cited (with the exception of Erickson et al., 2004) did not use additional selection mechanisms in their media, and from their results, it is not clear whether background organisms are included in the counts or not. It can be advised for future inoculation experiments with BSFL, with authorized or not (yet?) authorized substrates, to anticipate on the abundance of organisms closely related to the wildtype target organism and include an extra selective or elective aid.

In our experimental set-up, we incorporated two types of uninoculated conditions, being the substrate alone and the substrate containing larvae. In the first two experiment series, we discovered that the uninoculated samples did not remain free of *Salmonella*. This was true for the two series and for both substrate and larvae. This finding, and taking into account our precautions to avoid cross-contamination during manipulations, urged us to conclude that the infection must have happened via transmission in the air (with 60% RH) in the ventilated climate chamber. This was confirmed by the fact that cross contamination did not occur when incubation of uninoculated and inoculated containers were incubated separately. While airborne transmission is generally not associated with *Salmonella* as a route for spreading in the food industry, multiple reports document on airborne transmission of *Salmonella* between animals in poultry (Adell et al., 2014; Gast et al., 1998; Holt et al., 1998; Kallapura et al., 2014; Lever & Williams, 1996; Richardson et al., 2003) and pig (Ikeguchi et al., 2005; Oliveira et al., 2006) houses. In a previous study by our research group on the dynamics of *Salmonella* sp. in mealworm rearing (Wynants et al., 2019), uninoculated and inoculates replicates were incubated together (in containers that were not covered and that were placed at the same distance from each other as in this study and using in the same climate chamber (at 28 °C and 65% RH) as in this study), and there was no cross-contamination. The mealworms were reared in wheat bran (without water addition), however, which is expected to have a much lower moisture content than the moistened chicken feed in the current study. It is assumed that the moistened substrate in this study facilitates airborne transmission, viability during transmission and growth upon arrival on a new location (e.g. a neighboring container) of *Salmonella* cells. It is reasonable to extrapolate these findings and possible explanations to large scale BSFL rearing in stacked, open crates in a production facility with moistened and circulated air. If one crate contains substrate and crawling larvae that are highly contaminated, a rapid spread to other containers via the air throughout the rearing facility may take place. Even when harvested larvae are further processed with treatments that may reduce or eradicate *Salmonella*, a massive outbreak in the rearing facility should be avoided.

## 5. Conclusions

Our study revealed that, when reared on chicken feed, BSFL does not show a reducing effect on *Salmonella* counts in the substrate. It can be concluded though, that outgrowth of *Salmonella* is slower when the initial contamination level is lower. In addition, our study demonstrates that airborne transmission is possible in laboratory conditions and we expect that it also may occur in industrial production facilities. Altogether, these observations lead to the general recommendation for insect producers to use substrate ingredients free of *Salmonella*, to avoid the entrance of the pathogen in the rearing and post-harvest processing line by any other route, and not to count on the antimicrobial activities that BSFL exert in some situations to eradicate the food safety risk. Future inoculation experiments are needed, using other *Salmonella* serotypes and other zoonotic bacteria, other substrates, larvae of other ages and variations on the rearing protocols to further elaborate on the dynamics of this pathogen and to support risk assessments. From our study, we can advise on the use of antibiotic resistant organisms to allow a proper monitoring of the inoculated strain. PCR technology can also assist in pathogen monitoring, provided proper primers for the target organism are available and background interference of the matrix can be excluded.

## 6. Acknowledgements

DV is financed by the Research Foundation - Flanders (FWO) via the SBO project ENTOBIOTA (S008519N) as well as by the European Union’s Horizon 2020 Research and Innovation program via the H2020 project SUSINCHAIN (grant agreement number 861976). LVM and DL are funded by the former and latter project, respectively. JDS holds a postdoctoral fellowship grant (grant number 12V5219N) of the FWO. We would like to thank RADIUS for providing the 8-day old BSF larvae to start the rearing experiments and Dr. Aaron White for sharing his plasmids to generate Kanamycine resistant *Salmonella* strains.

---

Supplementary Figure 1: Impact of *Salmonella* presence on larval growth in experiment series 1.

Supplementary Figure 2: Impact of selective medium containing kanamycin on growth of kanamycin-resistant *Salmonella* strains and background micro-organisms. A) Plating of sample extracted from larvae in design 1 (challenged with 7 log CFU/g *Salmonella*) on regular RAPID’*Salmonella* agar; B) Plating of sample extracted from larvae in design 2 (challenged with 7 log CFU/g Salmonella) on RAPID’*Salmonella* agar with kanamycin (50 μg/ml).

