## Supplementary figures and images for "Dynamics of *Salmonella* inoculated during rearing of black soldier fly larvae (*Hermetia illucens*) on chicken feed"

### Supplementary Figure 1

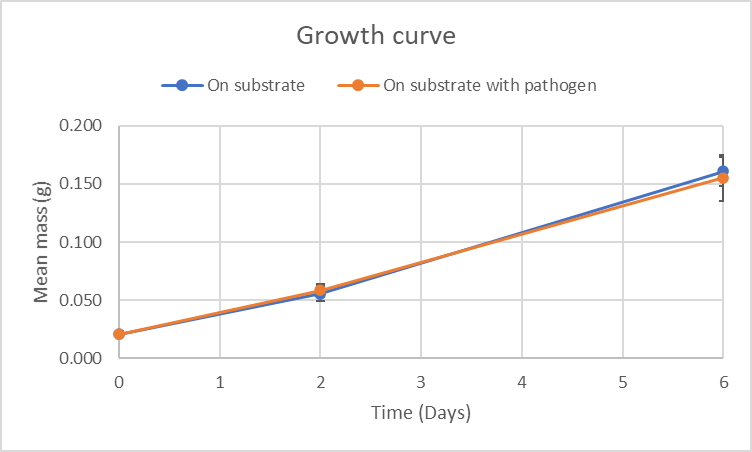

### Supplementary Figure 2

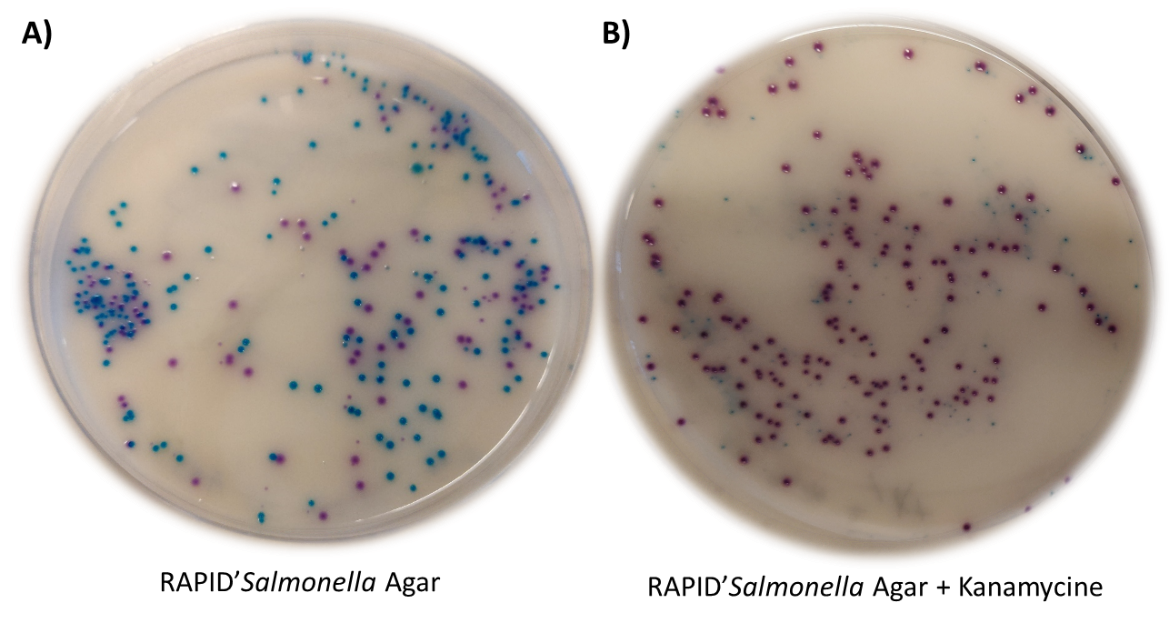
