## Supplementary Table 1 for "Dynamics of *Salmonella* inoculated during rearing of black soldier fly larvae (*Hermetia illucens*) on chicken feed"

**Supplementary Table 1:** PCR mixture and program for the confirmation of the incorporation of kanamycine resistance in *Salmonella*

| **PCR mixture** | | **PCR program** | | |
| --- | --- | --- | --- | --- |
| **Volume** | **Component** | **Time** | **Temperature** |  |
| 2 µl | Template (20-100 ng/µl) | 5 min | 95°C |  |
| 3 µl | 10x DreamTaq buffer | 45 sec | 95°C | Repeat 30 times |
| 0.1 µl | DreamTaq | 30 sec | 50°C |  |
| 0.6 µl | dNTPs (10 mM) | 1min30 | 72°C |  |
| 0.5 µl | Primer F (20µM) | 5 min | 72°C |  |
| 0.5 µl | Primer R (20µM) | ∞ | 12°C |  |
| 22.9 µl | millQ water |  |  |  |
